# ACIS, A Novel KepTide™, Binds to ACE-2 Receptor and Inhibits the Infection of SARS-CoV2 Virus *in vitro* in Primate Kidney Cells: Therapeutic Implications for COVID-19

**DOI:** 10.1101/2020.10.13.337584

**Authors:** Gunnar Gottschalk, James Keating, Kris Kessler, Chi-Hao Luan, Konstance Knox, Avik Roy

## Abstract

Coronavirus disease 2019 (COVID-19) is a severe acute respiratory syndrome (SARS) caused by a virus known as SARS-Coronavirus 2 (SARS-CoV2). Without a targeted-medicine, this disease has been causing a massive humanitarian crisis not only in terms of mortality, but also imposing a lasting damage to social life and economic progress of humankind. Therefore, an immediate therapeutic strategy needs to be intervened to mitigate this global crisis. Here, we report a novel KepTide™ (Knock-End Peptide) therapy that nullifies SARS-CoV2 infection. SARS-CoV2 employs its surface glycoprotein “spike” (S-glycoprotein) to interact with angiotensin converting enzyme-2 (ACE-2) receptor for its infection in host cells. Based on our *in-silico*-based homology modeling study validated with a recent X-ray crystallographic structure (PDB ID:6M0J), we have identified that a conserved motif of S-glycoprotein that intimately engages multiple hydrogen-bond (H-bond) interactions with ACE-2 enzyme. Accordingly, we designed a peptide, termed as ACIS (**<u>AC</u>**E-2 **<u>I</u>**nhibitory motif of **<u>S</u>**pike), that displayed significant affinity towards ACE-2 enzyme as confirmed by biochemical assays such as BLItz and fluorescence polarization assays. Interestingly, more than one biochemical modifications were adopted in ACIS in order to enhance the inhibitory action of ACIS and hence called as KEpTide™. Consequently, a monolayer invasion assay, plaque assay and dual immunofluorescence analysis further revealed that KEpTide™ efficiently mitigated the infection of SARS-CoV2 *in vitro* in VERO E6 cells. Finally, evaluating the relative abundance of ACIS in lungs and the potential side-effects *in vivo* in mice, our current study discovers a novel KepTide™ therapy that is safe, stable, and robust to attenuate the infection of SARS-CoV2 virus if administered intranasally.

## Introduction

COVID-19 is a viral disease that causes severe acute respiratory syndrome with significant mortality [1, 2]. This highly contagious disease is caused by a viral strain named as Coronavirus-2 (SARS-CoV2) [3]. A recent study [4] has shown that SARS-CoV2 employs its membrane-bound “S-glycoprotein” to interact with the host receptor ACE-2 and that interaction plays the most critical role in the entry of viral particles in human cells. However, the molecular details of that complex formation were not properly understood. Previously, Wu *et al* employed X-ray diffraction strategy to resolve the 3D structure of S-glycoprotein of Coronavirus-1 (SARS-CoV1) and ACE-2 at a resolution of 3Å [5]. According to the structural hallmarks [6], a single subunit of S-glycoprotein was shown to interact with ACE-2. However, a large heterotrimeric structural assembly of S-glycoprotein with ACE-2 was difficult to resolve, until recent report by Song et al [7]. Consistently, their Cryo-EM structural analysis revealed that S3 subunit of the heterotrimeric S-glycoprotein of SARS-CoV1 indeed employed its conserved motif for its interaction with ACE-2.

S-glycoprotein of SARS-CoV2 shares a significant sequence homology with that of SARS-CoV 1. Based on LALIGN pairwise-alignment study, SARS-CoV2 displays 92% sequence homology with Coronavirus-1 in S-glycoprotein. Consistently, with a SWISSMODEL homology modeling strategy, we identified a similar structural landmark in SARS-CoV2, which has been found to share equivalent interactions with ACE-2 protein of host. Therefore, we are interested to design a peptide from this motif that could bind to ACE-2, inhibit the interaction with S-glycoprotein and finally mitigate the infection of the virus in the host.

Peptide therapy is a front-line targeted therapy used for the treatment of diverse metabolic disorders. The application of this therapy has been very effective in the treatment of autoimmune disorders [8], neurodegenerative diseases [9, 10], cancer [11], and allergy [12]. However, its application in viral infection may not be effective primarily because viral infection is associated with the action of many serine and cysteine exoproteases[13] that can easily cleave cell surface proteins and peptides at low pH [14]. Here, we are introducing a new concept of peptide therapy known as KEpTide™ (Knock-End peptide) strategy, which is resistant to low-pH loving viral exoproteases [15] [16, 17]. In addition to that, one side of the peptide were modified with two chemical reactions that augmented basic or alkaline properties in both ends of the peptide. As a result, the peptide can never form a cyclic structure, which often causes the loss of function for small peptides [18]. In our present study, we designed two such ACIS KEpTide™, one is basic (Compound A-tagged KEpTide 1) and other is acidic in nature (Compound B-tagged KEpTide 2). Interestingly, our BLItz assay revealed that the Compound A-tagged basic KepTide™ (or **KEpTide 1**) displayed strong association with ACE-2 [Dissociation constant (K_D_) ~ 3nM]. Compound B-tagged acidic ACIS KepTide™ (or **KEpTide 2**) was also included in the study. A fluorescence polarization (FP)-based binding assay of that Compound B-tagged KepTide™ 2 also revealed a strong interaction with ACE-2. However, the dissociation constant (K_D_) (~ 6 nM) is almost two-fold higher than the K_*D*_ of Compound A-tagged KEpTide 1 suggesting that KEpTide 1 has stronger affinity to ACE-2 enzyme. Interestingly, a luminometric inhibition assay further indicated that both the KepTide™ displayed strong inhibition in binding of S-glycoprotein with ACE-2, however, again KEpTide 1 displayed stronger inhibition than KEpTide 2. Finally, KEpTide 1 was observed to strongly attenuate the infection of SARS-CoV2 in VEROE6 cells suggesting that our ACIS KepTide™ indeed has a strong therapeutic benefits in COVID-19.

## Results

### SARS-CoV1 and SARS-CoV2 share similar assembly with ACE-2

Recent study indicates that SARS-CoV2 engages receptor enzyme ACE2 of host to infect target cells. In fact, a Cryo-EM study [7] clearly identified that spike (S)-glycoprotein complex of SARS-CoV (Coronavirus-1) directly interacted with ACE-2. Therefore, neutralizing that interaction might have therapeutic prospect to prevent COVID-19 infection.

However, COVID-19 pandemic is caused by another strain of coronavirus named as SARS-CoV2 or coronavirus 2. Although, both coronavirus-1 and coronavirus 2 share almost 93% sequence homology in S-glycoprotein, it is not clear if SARS-CoV2 displays similar complex formation with ACE2 as because there was significant sequence disparity in the same structural hallmarks between two virus strains. To address this concern, we adopted a homology modeling analysis followed by a PyDock-based rigid-body docking strategy to resolve the complex formation of SARS-CoV2 and ACE-2 protein. To enhance the confidence of the prediction, we incorporated structural restraints in the same stretches of amino acids located in the target motif. Interestingly, we were able to derive similar structural outcome for SARS-CoV2 S-glycoprotein while forming complex with ACE2 enzyme (data not shown). We observed that a tridecapeptide in that conserved motif of SARS-CoV2 intimately engaged in interactions with ACE-2 protein.

### Designing ACIS peptide

Once, we determined the structural hallmark of the interaction between ACE-2 and S-glycoprotein, our next aim was to design a blocking peptide. Our designed peptide is a tridecapeptide, which was named as **<u>AC</u>**E-2 **<u>I</u>**nteracting motif of **<u>S</u>**-glycoprotein or ACIS peptide. To nullify the possibility of off-target effects, we searched the homology of this peptide against other host proteins with protein BLAST tool at NCBI server. Interestingly, the BLAST result followed by constraint-based multiple alignment analysis displayed no homology with any other protein except S-glycoprotein indicating that our ACIS peptide might exhibit a target-specific action of neutralizing the interaction between Coronavirus2 and ACE-2 enzyme.

To further confirm, next, we adopted a PyDock rigid-body structural analysis to verify if ACIS peptide could block the interaction between ACE-2 and S-glycoprotein. Strikingly, the peptide was found to have significantly impaired the interaction between ACE-2 and S-glycoprotein. In the presence of that peptide the entire S-glycoprotein was shifted far from the conserved β-sheet motif of ACE-2 enzyme. The peptide was predicted to form multiple strong H-bond interactions with ACE-2 enzyme nullifying the possibility to be outcompeted by S-glycoprotein (data not shown).

### Exploring physical interaction between ACIS and ACE-2

The major limitation of the peptide is its unpredictable secondary structure and the formation of a cyclic structure resulted by the reaction of amino and carboxy-terminal ends of peptide. For that reason, we modified one end of the peptide with two chemical reactions that nullified the possibility of forming cyclic structure. We termed these peptides as Knock-end peptide or KEpTide™. We generated two such KEpTides with two different compounds. Compound A-tagged ACIS was named as KEpTide1 and compound B-tagged second peptide was termed as KEpTide 2.

To understand the actual binding between KEpTides and ACE-2, we first performed BLItz label-free bio-layer interferometry assay that measures the binding of KEpTide 1 with ACE-2 as a function of incubation time. Interestingly, increasing doses (6 nM to 100 nM) of ACE-2 protein displayed a dose-dependent increase in binding with KEpTide1 (Fig. 1A) as indicated with increasing slope values of hyperbolic binding curves. The binding affinity was calculated based on the K_*D*_ values of binding curves derived from 15.3 nM and 30.6 nM of ACE-2. According to that analysis, KEpTide 1 displayed significantly strong binding with ACE-2 with K_*D*_ =2.86 nM (Fig. 1B).

**Fig. 1.**
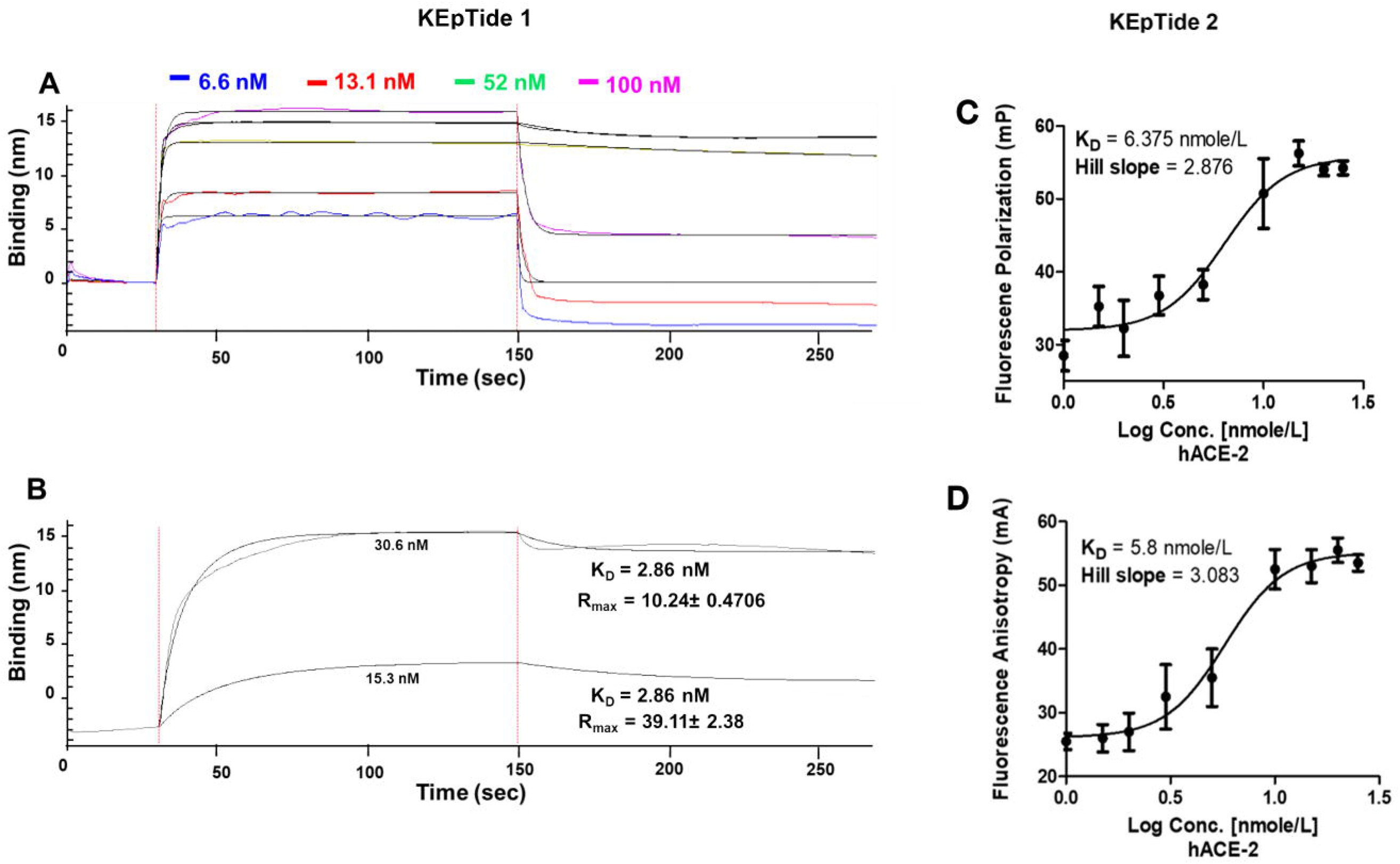
Biolayer Inferometry (BLItz)-based binding assay and Fluorescence Polarization (FP) assay to evaluate the interaction of KEpTides with hACE-2. (A) dose-dependent BLItz binding curve of hACE-2 (dissolved in buffer-1 of 20 mM HEPES, 150 mM NaCl, 0.05% Tween, pH 7.5) against a given concentration of KEpTide 1 [1.654 mg/ml (1mM) in DMSO and then dissolved in buffer-1 at a concentration of 2.5 μg/mL]. Increasing doses of ACE-2 starting from 6.6 nM to 100 nM were titrated with the given concentration of ACIS. Results were confirmed after 4 independent experiments. (B) A dose-response analyses to determine the accurate dissociation constant (K_D_ ~ 3 nM) and maximum response (R_max_) of binding between KEpTide 1 and hACE-2 (15.3 & 30.6 nM). Results were confirmed after 4 independent experiments. (C) dose-dependent FP binding curve of hACE-2 (dissolved in 20 mM HEPES, 150 mM NaCl, 0.05% Tween, pH 7.5) against a given concentration of KEpTide 2 (1.94 mg/mL in DMSO). Increasing doses of ACE-2 (1, 1.5, 2, 3, 5, 10, 15, 20, 25 nM) were titrated with the given concentration of ACIS (**K_D_ =6.38 nmole/L**). Results were confirmed after 4 independent experiments. (D) Fluorescence anisotropy data displays dose-dependent binding between increasing doses of hACE-2 and 1.94 mg/mL ACIS (**K_D_ =5.8 nmole/L**). Results were confirmed after 4 independent experiments.

To confirm the binding of ACIS KepTide™ with ACE-2, we performed another binding affinity assay known as fluorescence polarization. This assay measures the binding of KEpTide 2 with ACE-2 in the form of fluorescence potential (mP) or anisotropy (mA) of the depolarized fluorophore as a function of increased concentration of ACE-2. Interestingly, both fluorescence polarization (Fig. 1C) and anisotropy (Fig. 1D) analyses indicated that there was a strong binding between KEpTide 2 and ACE-2 with K_*D*_ value of approximately 6 nM. Interestingly, K_*D*_ value of KEpTide 1 (Fig. 1B; 2.86 nM) is two times more than that of KEpTide 2 (Fig. 1C; 6 nM) suggesting that KEpTide 1 displays stronger affinity towards ACE-2 protein. To nullify the off-target interaction of the ACIS KEpTide, we incubated 100 nM KEpTide 2 with an unrelated dehydrophos biosynthetic enzyme DHPH. Interestingly, 100 nM of peptide displayed significant binding only with ACE-2 (1 nM), but not with DHPH (5.5 nM) (Supplementary Fig. 1) suggesting that the binding of ACIS with ACE-2 is specific.

### Evaluating the inhibitory effect of ACIS KepTide™ on the interaction of S-glycoprotein with ACE-2

Next, we wanted to evaluate the if ACIS inhibited the interaction between S-glycoprotein and ACE-2. To demonstrate that, we performed a kit-based luminometric inhibitor screening assay that evaluated the efficacy of both KEpTide 1 and 2 in inhibiting the interaction between S-glycoprotein and ACE-2 protein. The assay was performed as described in manufacturer’s protocol. Interestingly, increasing doses of both KEpTides significantly inhibited the association of S-glycoprotein with ACE-2 as indicated with the IC_50_ values of 5.12 nM (Fig. 2A) for KEpTide 1 and 24.47 nM (Fig. 2B) for KEpTide 2. Moreover, the hillslope of the inhibition plot was significantly steeper in case of KEpTide 1 (−1.5) compared to KEpTide 2 (−2.4) suggesting that KEpTide 1 displayed stronger inhibition than KEpTide 2. In contrast to sigmoidal binding curve, there was no binding observed in case of scrambled KepTides. Taken together, this result suggests that KEpTide 1 significantly inhibits the interaction of S-glycoprotein with host ACE-2 protein and that inhibition is observed to be almost 5-times stronger than KEpTide 2.

**Fig. 2.**
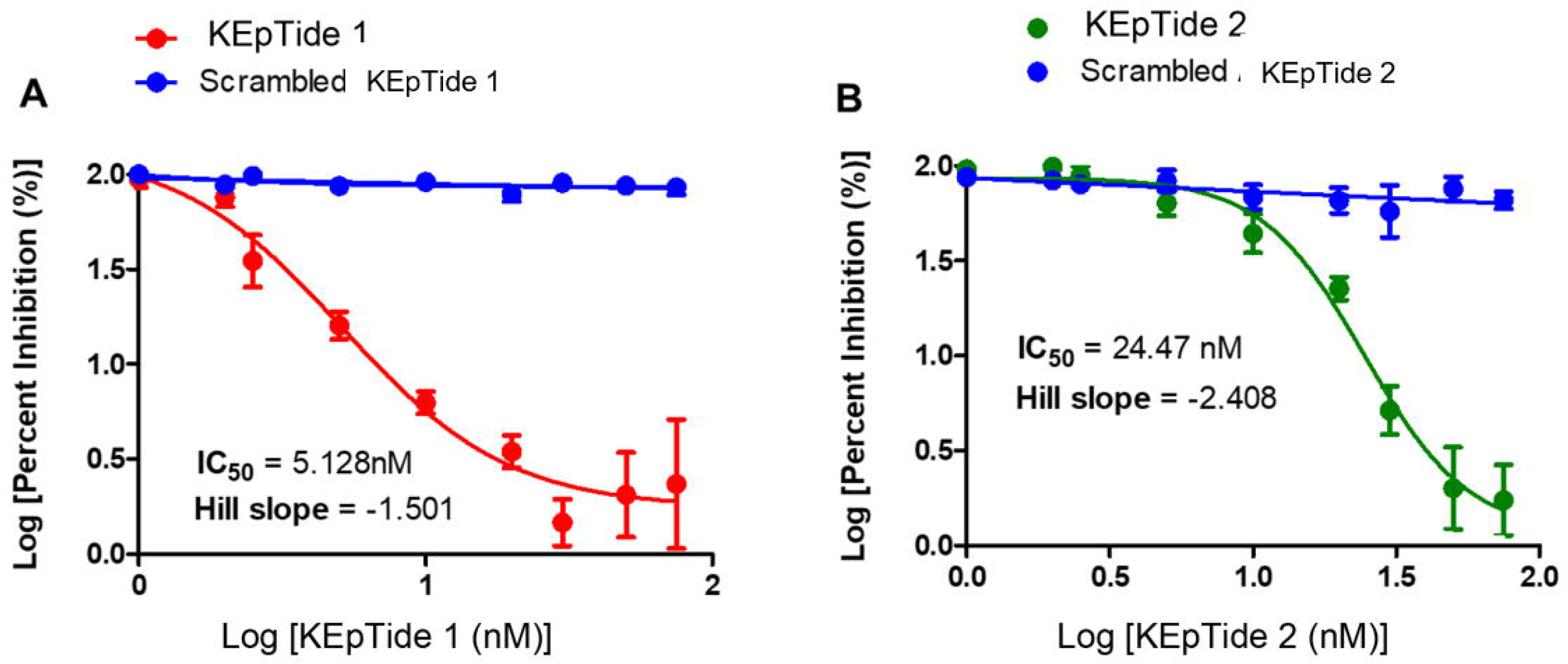
Effect of ACIS on inhibiting the complex formation between ACE-2 and S-Glycoprotein. Chemiluminometric binding assay of S-glycoprotein RBD and hACE-2(His-tag) with increasing doses of (**A**) KEpTide 1 (**B**) KEpTide 2 (0.5 nM to 100 nM) as directed with manufacturer’s protocol (BPS Bioscience, SARS-CoV2 inhibitor assay kit, Cat # 79936). IC50 value of KEpTide 1 (5.128nM) and KEpTide 2 (24.47 nM) were calculated after logarithmic value of percent inhibition as a function of log doses and then plotted with non-liner regression curve-fit analyses in GraphPad Prism software. Scrambled-peptide did not display any binding. Results are mean ± SD of three independent experiments.

### ACIS inhibits the infectivity of the SARS-CoV2 in VERO E6 cells

Until now we performed cell-free experiments to understand the effect of the KEpTide™ on ameliorating the interaction of S-glycoprotein with ACE-2. However, to delineate the effect of KepTides on nullifying the infectivity of the virus, cell culture experiments were needed. Therefore, next we did a series of cell culture study in mammalian VEROE6 cells. These cells are kidney cells of primate origin that strongly express ACE-2 receptor. Briefly, Wuhan standard stock of SARS-CoV2 (SARS CoV-2 USA_WA1/2020) was maintained in FDA-approved Coppe laboratories, enriched, titrated and applied on VERO (5*10^5^ cells per well with 90% confluency) cells at a dose of 1-2 PFU for the infection as described in a method section. In KEpTide-treatment condition, VERO cells were pre-incubated with 25 μM of KEpTide1 and 2 under serum-free condition for 30 minutes followed by the treatment with SARS-CoV2 strain for 2 and 6 hrs. In DMSO control group, VERO cells were treated with equivalent volume of DMSO. Two-hours timepoint was selected to evaluate the efficacy of the peptide to stop acute infection, whereas 6 hrs timepoint was selected for nullifying the secondary or chronic infection. After 2 hrs of viral incubation, supernatants with infectious virions were harvested from the top of the VERO cells and analyzed from agar monolayer invasion assay as described in method section. Briefly, different dilutions (1:20 to 1:800,000 of 5 *10^5^ virions) of viral supernatants were applied on agar-coated 96-well plate, neutralized with 10% neutral buffer formalin, and then stained with 2% Crystal Violet (CV) solution in 20% methanol. Interestingly, viral supernatants of 25 μM KEpTides-treated VERO cells, but not DMSO-treated cells, displayed significant damage in agar monolayer (Fig. 3A) as demonstrated in the loss of CV staining at 1:20 and 1:80 dilution series. Supernatants from both KepTides-treated VERO cells seemed to generate equivalent damage to the agar monolayer. However, further densitometric quantification of CV staining (Fig. 3A *lower panel*) demonstrated that there were almost 5-fold loss of agar monolayer in KepTides-treated groups compared to DMSO-treated VERO cells with partially stronger degeneration in KEpTide 1. These results suggest that KEpTide-treatment significantly attenuated the entry of SARS-CoV2 viral particles in the VERO cells causing their accumulation in the supernatants.

**Fig. 3.**
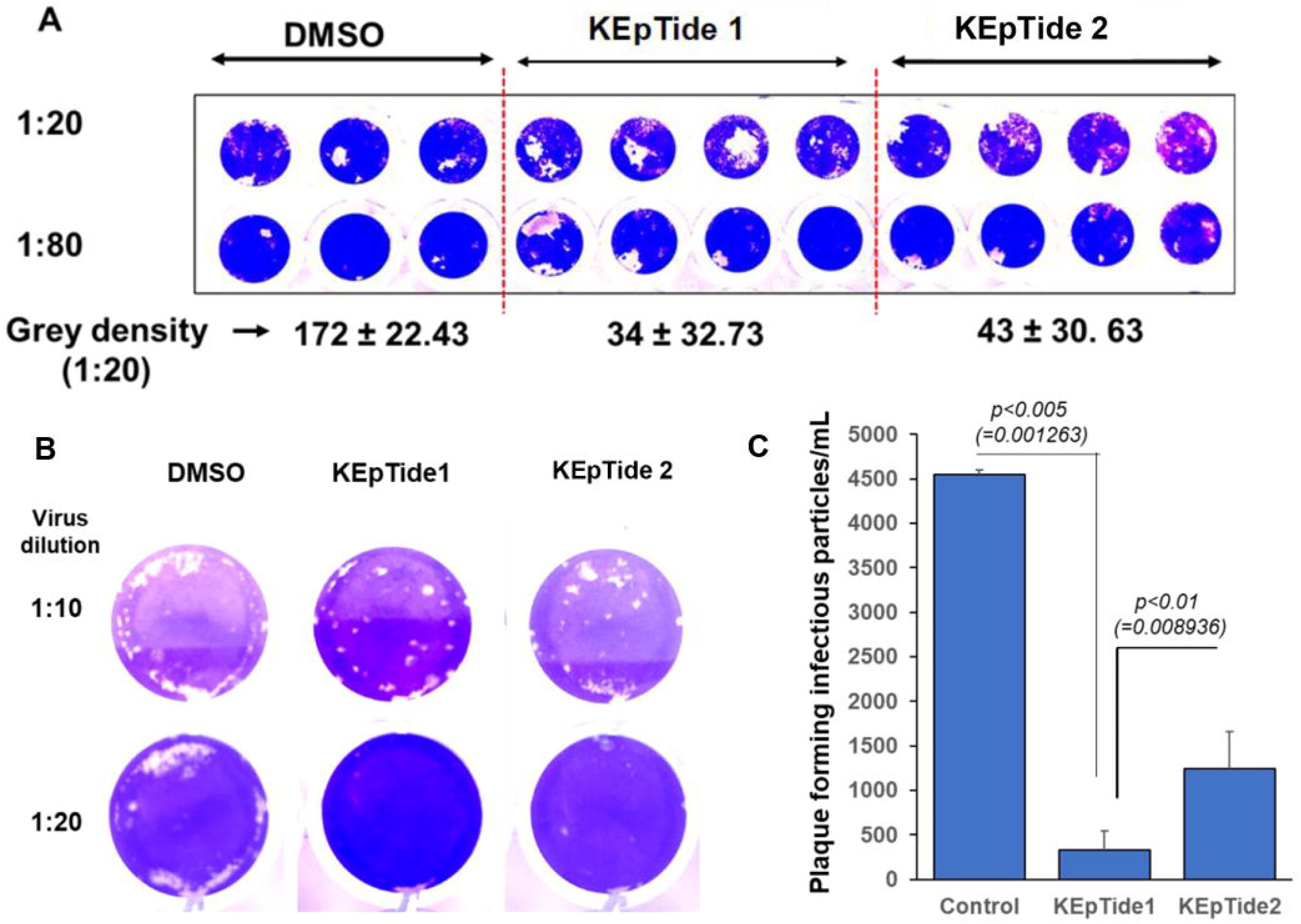
Cytopathic Effect Test and Plaque Assay to measure infectivity of residual virus present in the supernatants of KEpTide-treated VERO cells. **(A)** 5 × 10^6^ VERO cells were treated with 25 μM ACIS KepTides for 30 mins followed by the treatment of 1-2 MOI SARS-CoV2 (Stock# SARS CoV-2 USA_WA1/2020) for 2 hrs. After that, supernatants were harvested and then CPE test was performed as described in method section. Two dilution series of 1:20 and 1:80 was shown. Images were turned into greyscale followed by measuring the mean density in ImageJ software. (**B**) Ninety-percent confluent VERO E6 cells were treated with 25 μM KEpTide 1 and 2 followed by the treatment with SARS-CoV2 after different dilution starting from 1:10 to 1:800,000 (initial PFU =5 × 10^5^) as described under method section. After 72 hrs of virus infection, cells were overlaid with Agar and stained with 2% CV in ethanol. (**C**) Infectious particles were counted by counting numbers of single round plaque followed by normalizing with dilution factor. The final result was plotted as numbers of infectious particles per milliliter of inoculum. Results are mean ± SD of three independent experiments. The significance of mean was tested with One-way ANOVA with treatment as a single factor. The descriptive statistics results F_2,6_ =251.92 (>*F*_*c*_ =5.143); *p<0.0001*(= 1.63 × 10^−6^). The paired t-test was adopted to monitor the difference of means between control and KEpTide 1 as well as control and KEpTide 2.

Next, we performed plaque assay. To perform a plaque assay, 10- and 20-fold dilutions of a virus stock are prepared, and 0.1 ml aliquots are inoculated onto VERO cell monolayers. After 48 hrs of infection, VERO cells were covered with agar layer followed by staining with 2% CV. Each infectious particle generated a circular white zone indicative of infected cells whereas uninfected cells displayed strong blue color (Fig. 3B). Based on our densitometric quantification analyses, we observed that KEpTide 1 and 2 displayed significantly stronger protection, whereas KEpTide 1 displayed almost 15-fold stronger protection compared to control at 1:20 dilution dose (Fig. 3B & 3C). The similar statistics demonstrated only 3-fold protection in case of KEpTide 2.

To further confirm the profound activity of KepTides in the attenuation of viral infection, we performed a dual immunostaining analyses with ACIS KepTide (KEpTide2) (green) and S-glycoprotein (red). DAPI staining was carried out to identify big nuclei of VERO Cells. For this analyses, KEpTide 2 was preferred over KEpTide 1 as it was already labelled with fluorescence tag (green) and an excellent choice for double labelling with ACE-2 (red).Accordingly, after two hrs of SARS-CoV2 treatment (1-2 PFU), 25 μM of ACIS KepTide was observed to substantially inhibit the entry of viral particles (Fig. 4A), whereas no inhibition was observed in no DMSO, DMSO alone and scrambled-ACIS-treated control group. The result was further corroborated with a quantification study that clearly indicated a strong inhibition of viral load around ACIS-treated VERO cells (***p<0.0001) (Fig. 4B).

**Fig. 4.**
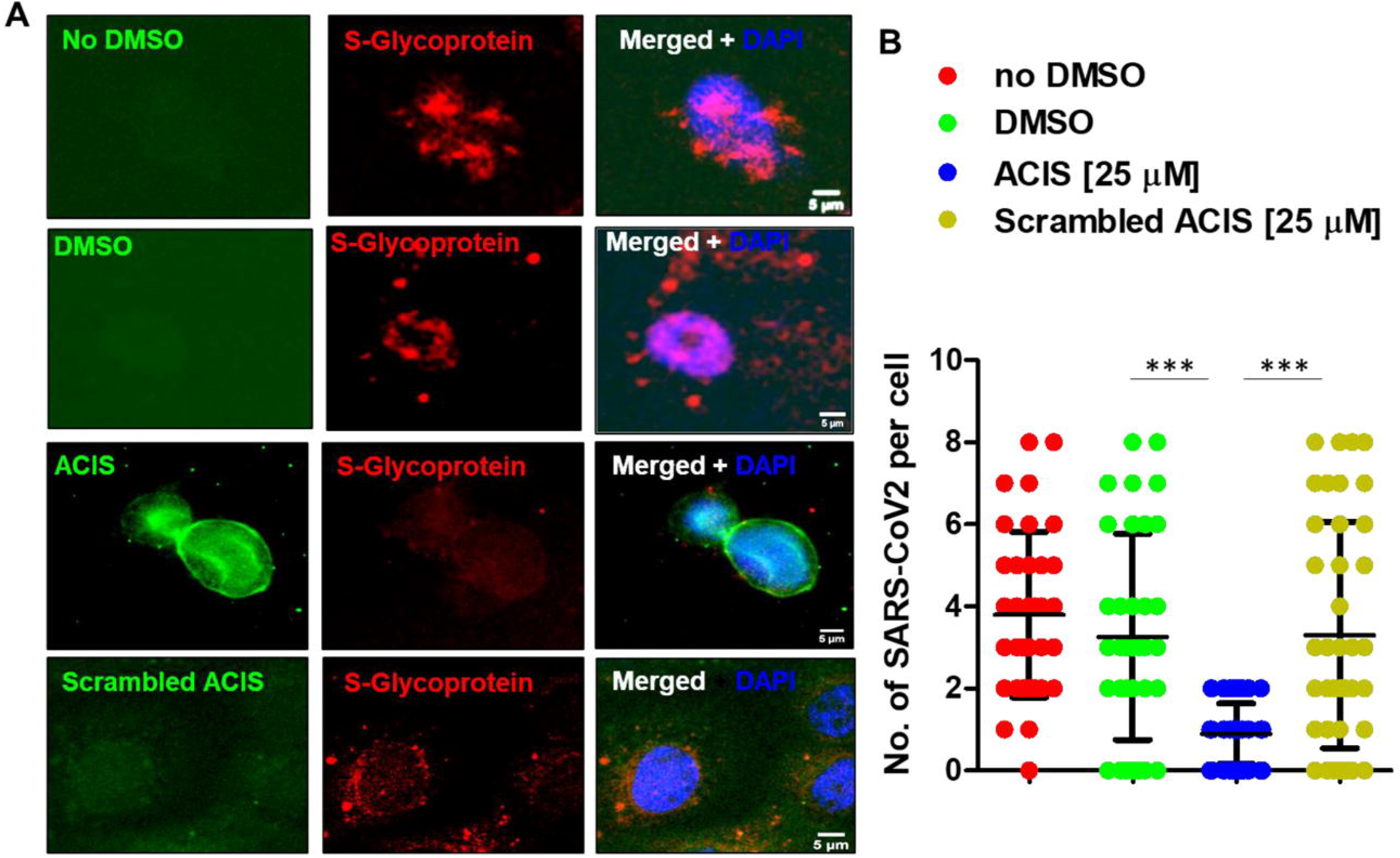
Effect of ACIS on the attenuation of acute infection (2 hrs) of SARS-CoV2 cells in VERO E6. **(A)** 5 × 10^5^ VERO cells were treated with 25 μM ACIS KepTides for 30 mins followed by the treatment of 1-2 PFU SARS-CoV2 for 2 hrs. After that, cells were fixed with 8% PFA and then dual immunolabeled with KEpTide 2 (KEpTide; green) and S-glycoprotein (red). (**B**) Forty VEROE6 cells per group were included to quantify number of S-glycoprotein-immunoreactive (*ir*) virus particles around VERO Cells. Nuclei were stained with DAPI. The significance of mean was tested with one-way ANOVA with treatment as an effector that results. *** F3_,156_ = 39.71 (>Fc =3.14); p<0.0001. Results are mean ± SD of three independent experiments.

In order to evaluate the efficacy of the KepTide in suppressing the secondary or chronic infection of SARS-COV2, next, we performed similar dual immunostaining procedure after 6 hrs of viral infection. At this time, virus was expected to initiate the second round of infection. Surprisingly, 25 μM of ACIS (KEpTide 2), but neither DMSO nor scrambled ACIS, continued to repel viral particles significantly from VERO cells (Fig. 5A) even at 6 hrs of infection. The effect was further confirmed with a quantitative analyses (Fig. 5B). Moreover, the distribution of KepTide molecule was carefully analyzed after 2 and 6hrs of viral incubation. We observed that ACIS was perfectly aligned on the membrane of VERO cells at 2 hrs, whereas significant numbers of KEpTide molecules were found to be internalized at 6 hrs suggesting that upon binding ACIS might also stimulate the internalization of ACE-2. As a result, ACIS KepTide efficiently inhibits the entry of SARS-CoV2 in host cells.

**Fig. 5.**
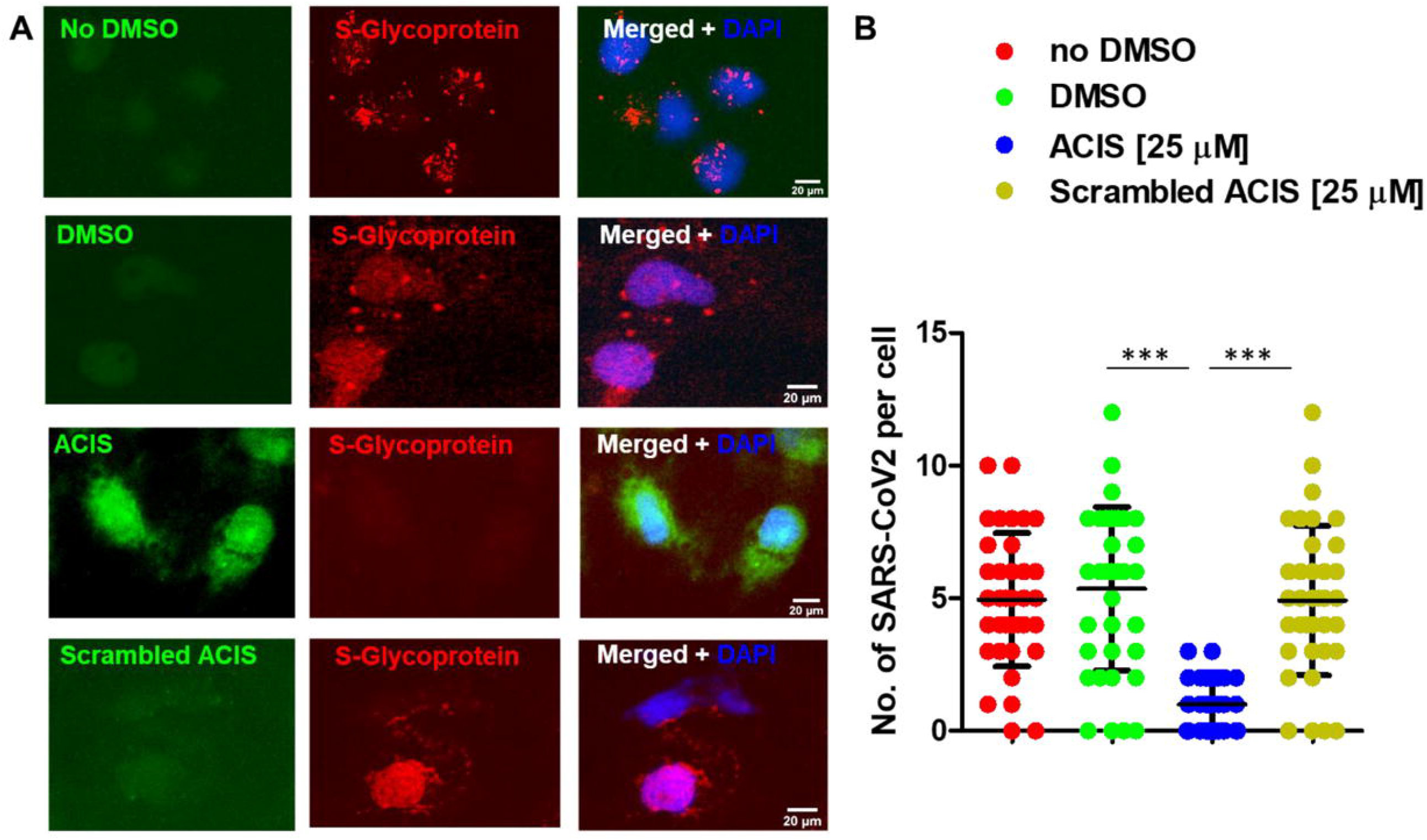
Effect of ACIS on the attenuation of secondary infection (6 hrs) of SARS-CoV2 cells in VERO E6. **(A)** 5 × 10^6^ VERO cells were treated with 25 μM ACIS KepTides for 30 mins followed by the treatment of 1-2 PFU SARS-CoV2 for 6 hrs. After that, cells were fixed with 8% PFA and then dual immunolabeled with KEpTide 2 (KEpTide; green) and S-glycoprotein (red). (B) Thirty-Forty VEROE6 cells per group were included to quantify number of S-glycoprotein-immunoreactive (*ir*) virus particles around VERO Cells. Nuclei were stained with DAPI. The significance of mean was tested with one-way ANOVA with treatment as an effector that results. The significance of mean was tested with one-way ANOVA with treatment as an effector that results. ***F_3,143_ = 24.36 (>Fc =6.19); p<0.0001. Results are mean ± SD of three independent experiments.

### Exploring half-life of KEpTide1 in Lungs after intranasal administration of KEpTide

So far, we have tested the efficacy of ACIS KEpTide on the attenuation of SARS-CoV2 infection *in vitro* in primate Kidney Cells. However, it is not known the effect of ACIS *in vivo*. Therefore, first we wanted to monitor how long the KEpTide can sustain in the target tissue once administered through intranasal route.

Therefore, next we wanted to assess the bioavailability of KEpTide1 in lungs and blood after administration of KEpTide1. Briefly, 0.1 mg/Kg bodyweight dose of KEpTide1 was administered intranasally for 0, 0.5, 1,2,6,12 and 24 hrs of time (n=6). After each time point, mice were sacrificed, and their blood and lung tissue were collected and weighed. For complete disintegration and compound A separation, lung tissue was homogenized with trypsin-containing PBS (1:1) for 10 mins at 37°C. The colorimetric assay (Fig.6) revealed that the intranasal administration of KEpTide1 increased its bioavailability with increasing time starting from 30 mins to 6 hrs timepoint with maximum at 1hr. Interestingly, the level of compound A was stable even after 6 hrs of treatment suggesting that the affinity of KEpTide is very strong for lung tissue (Fig. 6A). Moreover, we did not see any free compound A in blood in all these timepoints (Fig. 6B) suggesting that the half-life of compound A in blood might be very short or significant amount of that compound might be washed out through blood in early timepoint before 30 mins. Therefore, in future, we need to perform the same assay in shorter timepoints.

**Fig. 6.**
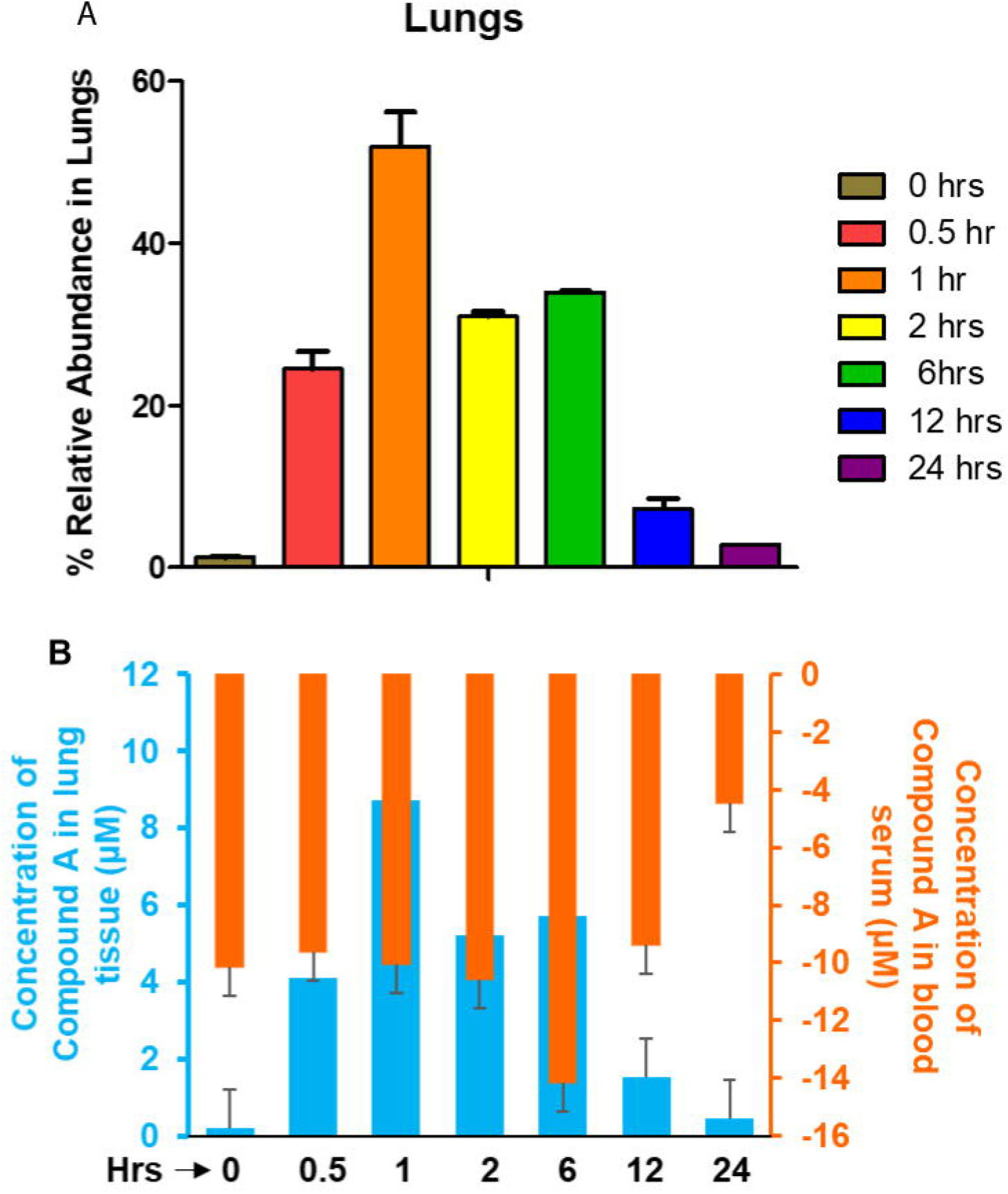
The assessment of the bioavailability of KEpTide 1 in lungs and blood. Eight to-10 weeks old BALB/C mice (n=6 with 3 males and 3 females) were given 0.1 mg/Kg bodyweight KEpTide1 at 0, 0.5, 1, 2, 6, 12 and 24 hrs. At each time point mice were sacrificed, lung tissue harvested, washed with 1X PBS, weighed, homogenized with 1:1 diluted trypsin (v/v with 1X PBS), centrifuged and then assayed for compound A as per manufacturer’s protocol. (A) Relative abundance (RA) was calculated considering the intranasal dose of KEpTide as 100% and plotted in a percent scale. (B) Absorbance values were converted to the absolute amount after fitting the absorbance value with the slope and intercept of standard curve having five different concentrations of 3.12, 6.25, 12.5, 25 and 50 μM of compound A standards. Results are mean ± SD of three different experiments. Significance of mean was tested with one-way ANOVA considering treatment as a single factor that results F6,14 =102.869 (>Fc = 2.84); *p <0.0001 (= 9.14 × 10^−11^).

### ACIS does not display any off-target side effects

It is possible that chronic administration of ACIS might generate some adversary off-target effects *in vivo*. To address that concern, next we monitored the effect of KEpTide *in vivo* in mouse over 10 days period following daily intranasal administration. We administered 0.1 mg /Kg body weight KEpTide1 intranasally to 8-10 weeks old BALB/C mice every day for 10 days. Each day, we monitored body weight, body temperature, heart rate, and oxygen saturation. Other parameters such as skin turgor, diarrhea and respiratory rate were monitored at Day 10. Interestingly, ACIS KEpTide 1 did not cause any loss in body weight (Fig. 7A), temperature (Fig. 7B), heart rate (Fig. 7C) and respiratory health (Fig. 7D) in these aged BALB/C animals suggesting that KEpTide 1 does not cause any toxic effects in vivo. Moreover, we did not see other health issue such as skin roughness or diarrhea in KEpTide1-treated BALB/C mice at the end of day 10. Although change in body weight showed significance (p<0.0008), but the result was confounded with two factors. *First*, the baseline value was low in KEpTide-treated animals. Across the treatment regime, KEpTide-treated animals did not gain any weight as evident from the comparison of baseline average weight (day 0) and the weight at the endpoint (day 10). The average weight remained same throughout KEpTide treatment. Second, the saline-treated mice gained significant weight. Therefore, taken together, these results suggest that KEpTide1 does not have any toxic off-target effects on health in terms of loss of body weight, hypothermia, hypertension rate and respiratory stress.

**Fig. 7.**
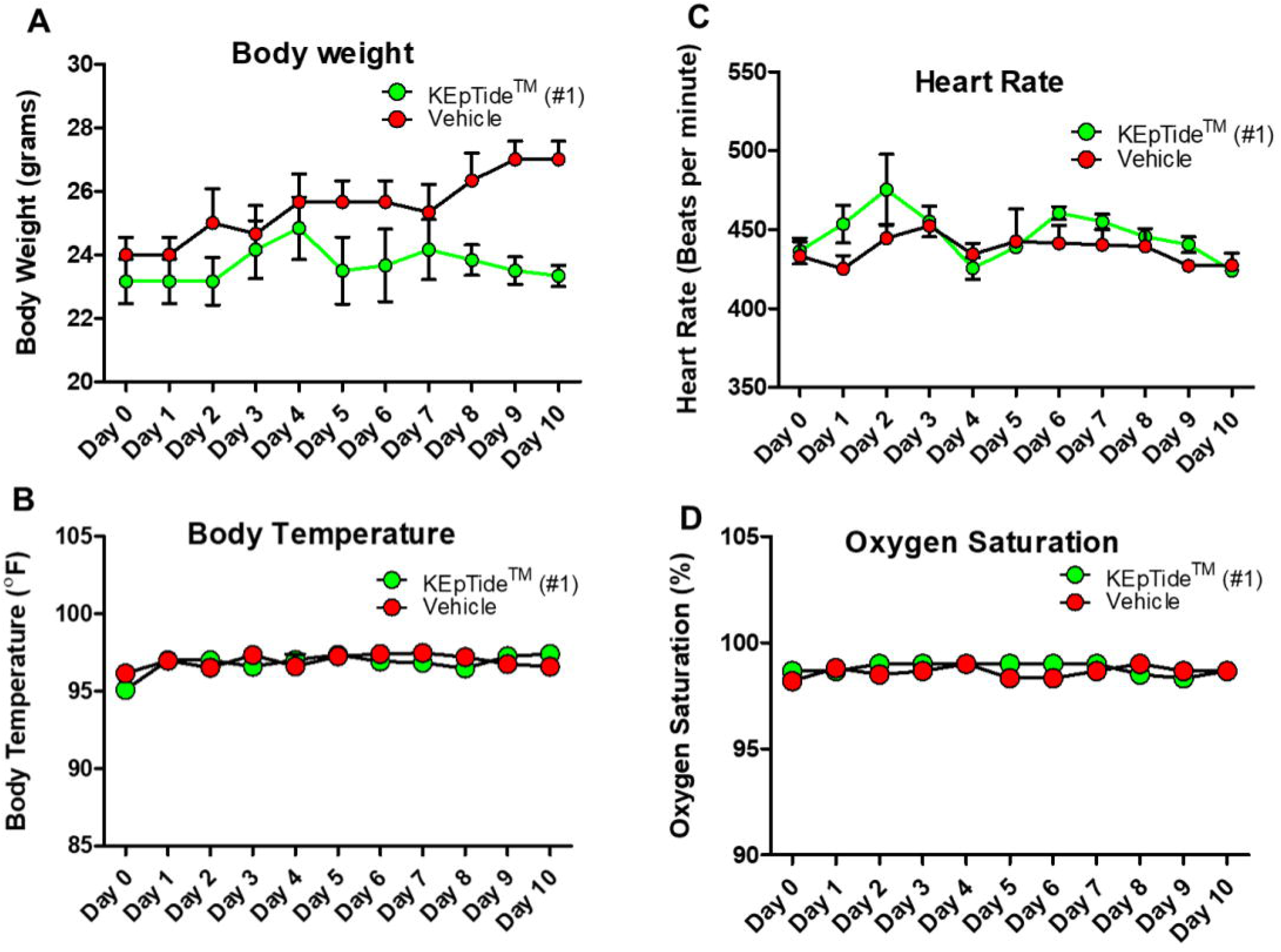
The effect of intranasally administered of KEpTide 1 on the health. (A) Eight-to-ten weeks old BALB/C mice were administered intranasally with KEpTide 1 at a dose of 0.1 mg/Kg bodyweight for 10 days (n=6 per group; 3 males and 3 females). The daily administration of KEpTide was performed in the afternoon, whereas the health issues were monitored in the morning. Different health parameters sch as (A) body weight, (B) body temperature, (C) heart rate, and (D) oxygen saturation was monitored starting from day 0 to day 10. Results are mean ± SD of three different experiments. Significance of mean between two groups were tested with paired t-test (p = 0.0008 for body weight; p = 0.5541 for body temperature; p = 0.0758 for heart rate and p = 0.3217 for oxygen saturation). Results are mean ± SEM of three independent experiments.

## Discussion

### Conceptualization the idea of KEpTide™

ACIS was designed from a conserved motif of S-glycoprotein and also observed to strongly impair the interaction between S-glycoprotein and ACE-2. Recently, a study performed by Goutret et al [19] has found that the repurposing of anti-malaria drug hydroxychloroquine (HCQ) significantly attenuated viral load in COVID-19 patients and this beneficial effect was further enhanced in combination with macrolide anti-bacterial drug such as azithromycin (AZT). However, until now, the trial of Goutret and colleagues does not provide sufficient evidence to support wide-scale application of HCQ treatment for the treatment of COVID-19, mainly due to the lack of rigorous methodology, cohort procedure and analysis. Moreover, HCQ treatment itself has several adverse side-effects including severe abdominal pain, fatigue, depression, hair loss, irregular heartbeat, and cardiac failure. In combination with strong antibiotic like AZT, HCQ might impose severe comorbidity. Moreover, in response to non-specific action of HCQ, there always is a risk for the generation of HCQ-resistant strain of coronavirus that might impose even higher threat to humankind. Repurposing of other anti-retroviral drugs such as Ebola drug remedesivir [20] and HIV drugs including lopinavir and ritonavir [21] are reported to display protective effect against COVID-19. However, these drugs are associated with severe side effects in COVID-19 patients such as respiratory failure, depigmentation of skin and severe anemia. Considering all these facts together, the current illness demands a specific therapeutic strategy that would only cure COVID-19 disease with less or no side-effect. Our ACIS peptide is the answer of that much-awaited targeted therapy. It is specifically designed from the part of S-glycoprotein that is conserved only in the coronaviral family therefore nullifies the possibility of any off-target effect. In fact, our *in vivo* experiments revealed that intranasal administration of KEpTide 1 did not cause any loss in body weight, hypothermia, hypertension, respiratory stress, diarrhea, and loss of skin tone.

BLAST analysis further confirmed that the peptide did not share any homology with other endogenous protein nullifying the possibility of non-specific action of the peptide. Interestingly, ACIS showed strong affinity towards ACE-2 only, not with other proteins. We tested the binding of ACIS with another common membrane protein DHPH and we did not see any affinity suggesting that ACIS is specific for only ACE-2. To further confirm the specificity of ACIS peptide towards ACE-2 protein, our BLItz assay between compound A-tagged ACIS and full-length ACE-2 revealed that there was a strong affinity of that peptide towards ACE-2 protein with K_D_ =3 nM. Interestingly, when acidic end of ACIS was modified with compound B, we saw almost two-times less affinity with ACE-2 and five-times less efficacy in inhibiting the interaction between ACE-2 and spike. These observations intrigued us to explore the chemistry of two KEpTides. Compound A-based modification increased the basicity of the KEpTide 1, whereas Compound B-based modification did not add any basicity to the KEpTides suggesting that the augmentation of basic property in the KEpTide enhanced its inhibitory effect on the binding of S-glycoprotein with ACE-2.

### Does KEpTide inhibit the infection of SARS-CoV2 virus?

Based on our monolayer invasion assay and plaque assay, we confirmed that both KEpTides significantly inhibited the entry of SARS-CoV2 virus in non-human primate cells. Quantification of infectivity of these virus displayed almost 15 fold more protection of entry in KEpTide 1 treated cells. To understand the mechanism of inhibition, we performed a dual immunostaining of KEpTide and S-Glycoprotein that displayed a strong membrane localization of KEpTide and subsequent repulsion of viral particles from VEROE6 cells. Our immunofluorescence results indeed demonstrated that even after 6 hrs of KEpTide-treatment completely inhibited the entry of the actual Wuhan standard virus in VEROE6 cells. Modification with heterocyclic compound A facilitates the nasal absorption of therapeutic agents. Therefore, next we monitored how efficiently that KEpTide1 can be administered through nasal route. Interestingly, intranasal administration of KEpTide1 efficiently delivered and maintained significantly high-level of KEpTide 1 in lungs even after 6 hrs suggesting that the intranasal administration of KEpTide efficiently delivered the therapy to the lungs. Moreover, our toxicity assay further demonstrated that long-term administration of KEpTide 1 did not cause any adverse health issue such as loss of body temperature, increased heart rate, loss of body weight and respiratory stress.

### Does KEpTide inhibit the endogenous action of biological ligands of ACE-2

Biological action of ACE-2 enzyme has been well studied. ACE-2 degrades angiotensin-II, a hormone that causes vasoconstriction. Therefore, activation of ACE-2 might stimulate dilation of blood vessel and lowing blood pressure. Endogenous peptide hormone angiotensin II [22], growth factors collectrin [23], calmodulin [24], and small sugar moiety N-glucosamine bind to the ectodomain of ACE-2. These interactions are not only required for the cellular metabolism of peptide hormones, but also required to retain the catalytic action of ACE-2 in plasma membrane. Interestingly, that same ectodomain plays critical role in the interaction with S-glycoprotein of SARS-CoV2. Hence, if a peptide is designed from the ectodomain of ACE-2, then that peptide binds to these biological regulators of ACE-2. Eventually, the catalytic action of ACE-2 will be disrupted. On the other hand, ACIS peptide is designed not from the enzyme, but against viral protein. Therefore, the peptide is only expected to inhibit the binding of exogenous SARS-CoV2 with ACE-2 enzyme and does not impair the interaction of other endogenous regulators with ACE-2.

### ACE-2 is a peptidase. Can it cleave ACIS peptide?

ACE-2 is a carboxypeptidase that cleaves carboxy-terminal peptide bond between proline and phenylalanine of angiotensin II. Our peptide does not provide that substrate specificity to ACE-2 mitigating the possibility of degradation of ACIS by ACE-2 enzyme. In addition to that, the basic nature of the KEpTide 1 further makes our peptide resistant to be cleaved by low pH peptidase.

Overall, our current report highlights a robust and targeted therapy against COVID-19 that can mitigate viral infection if administered through nasal route. Additional *in vivo* experiments are under progress that would demonstrate its prospect in human trial.

## Method

### Reagents and Materials

Compound A-tagged and Compound B-tagged ACIS KepTides were designed in house and commercially synthesized. S-glycoprotein antibody (Anti-SARS Antibody, S Specific, clone 154C; Cat # MAB8783) was purchased from Millipore Sigma. Donkey anti-mouse Alexa 647-conjugated affinity pure secondary antibody (Cat # 715-605-150) was purchased from Jackson ImmunoResearch Laboratories Inc. Spike RBD (SARS-CoV-2): ACE2 Inhibitor Screening Assay Kit was purchased from BPS Biosciences (Cat # 79931). Experiments with SARS-CoV2 strain were performed in Coppe Laboratories (CDC-approved and CLEA-certified) under the supervision of Dr. Konstance Knox.

### Determining 3D structure of S-glycoprotein of COVID-19

To understand the structural details of S-glycoprotein, we had adopted in silico homology modeling strategy to build 3D structure of S-glycoprotein. Initial structure was modeled by Swiss-Model server, which is operated with a program known as Deep View 3.7β2, an online macromolecular analytical tool of Expert Protein Analytical System (ExPASy). The sequence of S-glycoprotein was derived in FASTA format from the sequence available in PubMed with accession number QIC53213.1 and GI number 1811294675. SwissModel generated 3D structure based on the homology of spike protein of coronavirus-1 (PDB ID: 3SCL). The quality of the modeled structure had been evaluated with Quality Measurement Analysis tool (QMEAN) score −2.25. QMEAN is a composite scoring tool that estimates the global quality of the entire model as well as the local per-residue analysis of different regions within a model. Residue-level interaction was evaluated by Cβ atom potential and long-range interactions were validated by all-atom potential. A solvation potential was implemented to analyze the burial status of the residues. The local geometry of the derived structure was analyzed by a torsion angle potential over three consecutive amino acids. Based on all these energy scores, the best predicted structure of S-glycoprotein was achieved.

### Molecular docking of ACE-2 protein with ACIS peptide

In order to understand the interaction between ACIS peptide and ACE-2 in a molecular level, Pydock, a rigid body protein-protein docking tool had been applied. The most stable docked-structure had been resolved based on electrostatic (Eele), desolvation (Edesolv) and Van der Waals (ΔGVDW) energies and finally displayed with Chimera software. According to that analysis, ACIS peptide had been found to be docked in the interface of Spike glycoprotein and ACE-2 protein.

### BLItz assay

The assay was performed in the BLItz platform (FortéBio) (Northwestern Keck biophysics facility) applying high precision streptavidin Biosensors (FortéBio). The biosensor was hydrated in a 96 well plate with the working buffer and then loaded in the machine. The lyophilized KEpTide 1 peptide was diluted in DMSO at a concentration of 2.5 μg/mL and then loaded in biosensor. After that, the KEpTide 1 was titrated with different concentrations of ACE-2 protein, which was diluted in a buffer containing 0.05% Tween-20, 20mM Hepes, 150 mM NaCl, pH7.5. The analysis was performed following the protocol provided by the BLItz Pro software.

### Fluorescence Polarization

Compound B-labeled KEpTide 2 was diluted in DMSO at a concentration of 2.5 μg/mL and 100 nM final concentration of KEpTide 2 were mixed in protein dilution buffer (20 mM Hepes,150mM NaCl pH7.5). A black-wall 384-well microplate (Corning, Lowell, MA) was loaded with a serially diluted compound B-tagged ACIS and then reacted with a 2μM fixed concentration of ACE-2 protein. The polarization potential (mP) and anisotropy (mA) readings were obtained using a Tecan microplate reader (Tecan) at the Structural Biology Facility at Northwestern University. All curve fitting and data analysis were performed using Igor Pro (Wave Metrics, Portland, OR). Curve-fitting was done in GraphPad Prism 8 software using log[inhibitor] versus response, variable slope liner regression curve analysis module and resultant IC_50_ value was calculated.

### Inhibitor screening assay

SARS-CoV2 inhibitor screening assay was performed with the luminometric detection method as mentioned in the manufacturer’s protocol (Cat# 79931; BPS bioscience).

### Measurement of Cytopathic Effect (CPE) of SARS-COV2 from the supernatant of ACIS-treated VERO Cells

For making sufficient viral stock, 90% confluent VERO cells were infected with 5 × 10^5^ PFU SARS-CoV2 Wuhan standard (SARS CoV-2 USA_WA1/2020) at a MOI of 0.025. VERO cells were grown in Growth in complete DMEM media supplemented with 10% FBS. Before infection, the media was changed to complete media supplemented with 2% heat inactivated FBS. After 72 hrs of infection, viral stock (Media of T75 flask) was harvested, and RT-PCR was run to quantify the genomic equivalent. To test the effect of ACIS peptide on the CPE of viral particles, 5X 10% VERO cells were treated with 25 μM of ACIS or equivalent DMSO for 30 mins under serum-free condition. After that, unbound ACIS was aspirated followed by the infection with 1-2 genomic equivalent of SARS-CoV2 for 2 hrs. After 2 hrs, supernatants were harvested and applied in an agar monolayer-coated 96-well plates at a different dilutions starting from 1:20 to 1:800,000. The supernatants were kept at 37°C for 72 hrs, aspirated and fixed with 10% formalin for 30 mins at room temperature. Once the formalin is aspirated, 2% Crystal Violet solution (v/v n 20% methanol) was added in each well for 5 mins, rinsed and dried. The CPE was monitored by counting the extent and numbers of holes in the agar monolayer.

### Plaque Assay

SARS-CoV2 infectious titer was analyzed by plaque assay according to a previous report [25, 26] with slight modifications. Briefly, VERO cells were split into 6-well culture plates at the concentration of 5 × 10^5^ cells/well. An aliquot of SARS-CoV2 stocks was thawed and then 100-fold serially diluted in the culture media starting from 1:20 fold dilution. VERO E6 cells were pre-incubated with DMSO, 25 μM of KEpTide 1 and 2 for 30 mins and then inoculated with 500 μL of each SARS-CoV2 dilution and incubated at 37 °C for 1 h with rocking every 15 min. The inoculum was removed, and the cells were washed once with 1X PBS to eliminate the unbound virus particles. The cells were overlaid with overlay media containing 1.25 μg/mL acetylated trypsin and 0.8% (w/v) agarose (Lonza)]and additionally incubated at 37 °C for 72 h. Cells were fixed and stained in 20% ethanol containing 2 % crystal violet at room temperature for 15 min to visualize the plaques. The infectious particles were counted per well, normalized with the dilution factor, and counted as a number of infected particles per mL of inoculum. The data represent the mean ± standard error of four independent experiments.

### Double Immunocytochemistry

For Immunocytochemical analyses, 5 × 10^6^ VERO Cells were plated per well in 8-well chamber slide. Cells were starved with serum-free DMEM media for 30 mins followed by the treatment with 25 μM compound KepTide 2. After another 30 mins, unbound KepTides were aspirated and then added SARS-CoV2 at a dose of 1-2 PFU/ cell. After 2 and 6 hrs of virus treatment, viral cells were removed, VERO cells were fixed with 8% PFA and kept at 4°C for overnight. Next day, cells were washed with 1 X PBS followed by blocking with 2% horse serum, incubation with primary antibody (1:500 dilution with 1 X PBS-tween) for 2 hrs at 37°C, washed with 1 X PBS, incubated with 2° antibody (1:250 dilution at PBST combined with 1% horse serum). After that, cells were washed with PBS three times with DAPI (1:10000 dilution in water) at final wash. Slide was covered with coverslip and then dried at room temperature at dark. Next day, the cells were imaged in BX51 Olympus microscope at UIC RRC core facility.

### Bioavailability assay of KEpTide

Eight to ten weeks-old BALB/C mouse were administered with KEpTide 1 at a dose of 0.1 mg/Kg body weight intranasally. Briefly, mice were anesthetized with Ketamine-Xylazine mixture followed by administration of 2 μL of KEpTide1 in each nostril. The KEpTide 1 was inoculated for 0 min, 30 mins, 1, 2, 6, 12 and 24 hrs (*n=3* per group). After that, mice were sacrificed, their lung tissue and blood harvested to evaluate KEptide in blood and lungs. Lung tissue was disintegrated with 1xPBS: Trypsin (1:1) solution for 10 mins in 37°C followed by homogenization and centrifugation at 12000 rpm for 5 mins. That method completely digested proteins and peptides leaving heterocyclic compound A untagged and available in the homogenate. Blood serum was collected in EDTA tube. Twenty μL of serum and tissue lysate was assessed for free compound A as per manufacture’s protocol of a colorimetric assessment. The assay was performed and recorded in a 96-well plate format of EPOCH Biotek plate reader. The endpoint absorbance wavelength was 500 nm, and the final absorbance was recorded in Gen5 software version 2.05.

### Intranasal administration of KEpTide and monitoring health-related side-effects in BALB/C Mice after intranasal administration of ACIS KEpTide 1

Eight to ten weeks-old BALB/C mouse were administered with KEpTide 1 at a dose of 0.1 mg/Kg body weight. A single dose of KEpTide 1 and saline (control) was administered through intranasal route (2.5 μL each nostril = total 5 μL) each day for 7 days (*n=6 per group; 3 male and 3 female*). For controlled intranasal delivery of drug, each time micropipette was loaded with 2.5 μL of KEpTide and dispensed through nasal cavity by firmly holding mouse in the palm. Every day, mice were monitored for their overall health such as body weight. temperature, heart rate, oxygen saturation, skin tone and diarrhea. The body weight was monitored with digital scale (CAMRY), whereas body temperature, heart rate and oxygen saturation were monitored with WENSUIJIA Vet health monitor.

## Acknowledgements and Author Contribution

The study is funded by Sotira seed fund managed by JK and KK. GG, AR, CHL, and KKNOX designed, performed, and analyzed the data. AR conceived the idea and wrote the manuscript. For business-related correspondence, contact JK.

