## Supplemental Figure for "ACIS, A Novel KepTide™, Binds to ACE-2 Receptor and Inhibits the Infection of SARS-CoV2 Virus *in vitro* in Primate Kidney Cells: Therapeutic Implications for COVID-19"


Running title: ACIS KepTide$^{TM}$ and COVID-19


To whom correspondence should be addressed:
Avik Roy, Ph.D.
SOTIRA LLC

Ph: (773) 263 8182

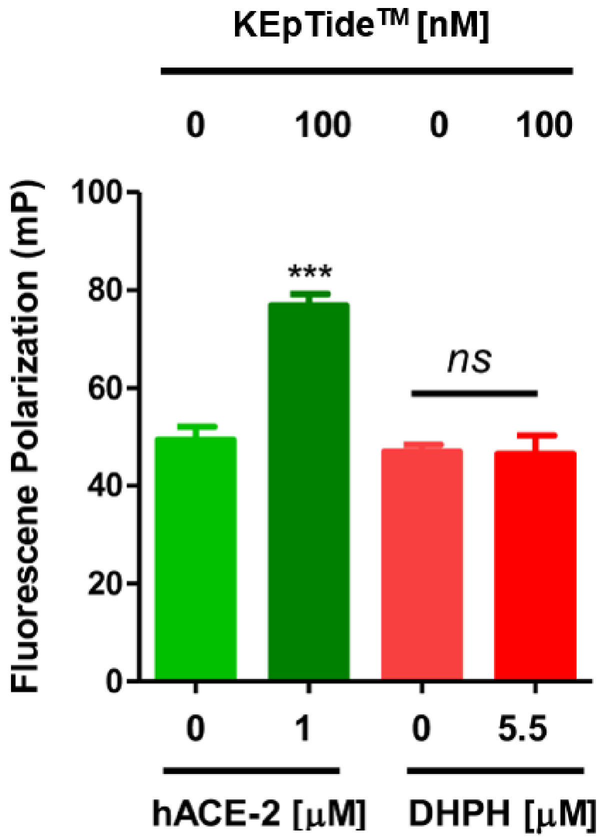


**Supplementary Fig 1. The Binding of ACIS with ACE-2 is specific.** FP analysis with 100 nM KEpTide 2 reacted with 1 nM of ACE-2 (green bars) and 5.5 nM of dehydrophos biosynthetic enzyme DhpH (red bars). Results are mean ± SD of three independent experiments. ***$p<0.0001$ versus control as tested with paired t-test; NS = no significance.
